# Simultaneous quantification of protein-DNA contacts and transcriptomes in single cells

**DOI:** 10.1101/529388

**Authors:** Koos Rooijers, Corina M. Markodimitraki, Franka J. Rang, Sandra S. de Vries, Alex Chialastri, Kim de Luca, Dylan Mooijman, Siddharth S. Dey, Jop Kind

**Affiliations:** Oncode Institute, Hubrecht Institute–KNAW and University Medical Center Utrecht, Utrecht, The Netherlands; Department of Chemical Engineering, University of California Santa Barbara, Santa Barbara, CA 93106, USA; Center for Bioengineering, University of California Santa Barbara, Santa Barbara, CA 93106, USA; Present address: Genome Biology Unit, European Molecular Biology Laboratory, Heidelberg, Germany

## Abstract

The epigenome plays a critical role in regulating gene expression in mammalian cells. However, understanding how cell-to-cell heterogeneity in the epigenome influences gene expression variability remains a major challenge. Here we report a novel method for simultaneous single-cell quantification of protein-DNA contacts with DamID and transcriptomics (scDamID&T). This method enables quantifying the impact of protein-DNA contacts on gene expression from the same cell. By profiling lamina-associated domains (LADs) in human cells, we reveal different dependencies between genome-nuclear lamina (NL) association and gene expression in single cells. In addition, we introduce the *E. coli* methyltransferase, Dam, as an *in vivo* marker of chromatin accessibility in single cells and show that scDamID&T can be utilized as a general technology to identify cell types *in silico* while simultaneously determining the underlying gene-regulatory landscape. With this strategy the effect of chromatin states, transcription factor binding, and genome organization on the acquisition of cell-type specific transcriptional programs can be quantified.

## Main

mRNA output is tightly regulated at many levels to ensure the precise coordination of cell-type specific gene expression programs. On the transcriptional level, packaging of DNA into chromatin can control access of transcriptional regulators to functional DNA elements like enhancers and promoters. Higher levels of organization that contribute to the regulation of gene expression involve the spatial segmentation of the genome into compartments with transcriptionally permissive or repressive gene regulatory activities. Failure to integrate and coordinate the multi-layered regulatory control of gene expression can result in developmental defects and the commencement of disease. To understand the regulation of gene expression it is key to dissect the direct relationships between epigenetic and transcriptomic heterogeneity. To this end, it is pivotal to develop techniques that enable simultaneous measurements of the epigenome together with the transcriptome from the same cell.

Recent advances in measuring genome architecture (HiC, DamID)^1-4^, chromatin accessibility (ATAC-seq and DNaseI-seq)^5-7^, DNA methylation (5mC)^8-10^, DNA hydroxymethylation (5hmC)^11^ and histone PTMs post-translational modifications (ChIP-seq)^12^ in single cells have enabled studies to characterize cell-to-cell heterogeneity at the gene-regulatory level. More recently, multiomics methods to study direct single-cell associations between genomic or epigenetic variations and transcriptional heterogeneity^13-16^ have provided the first methods to directly link upstream regulatory elements to transcriptional output from the same cell. Protein-DNA interactions play a critical role in regulating gene expression and therefore we have developed a new technology to simultaneously quantify these interactions in conjunction with transcriptomic measurements from the same cell without requiring physical separation of the nucleic acids.

DamID involves the fusion of the *E.coli* Dam adenine methyltransferase to a protein of interest, followed by the *in vivo* expression of the fusion protein to enable detection of protein-DNA interactions. For single-cell applications, a major advantage of the DamID method is that it minimizes biochemical losses arising from antibody-based pulldowns or degradation of genomic DNA (gDNA) that occurs in bisulfite-based methods. Further, as DamID is an *in vivo* method, protein-DNA interactions can be measured over varying time windows and can also be used to record cumulative protein-DNA interactions^17^. Currently, no methods exist to quantify protein-DNA interactions for an arbitrary protein-of-interest and transcriptomes in single cells. We therefore chose to benchmark scDamID&T and compare it to the previously reported single-cell DamID (scDamID) method where lamina-associated domains (LADs) were detected using a Dam-LmnB1 fusion protein^2^. Furthermore, we exploited the expression of untethered Dam to obtain DNA accessibility profiles simultaneously with transcriptome measurements and employed the scDamID&T technology to generate combined and allele-resolved single-cell measurements in hybrid mouse embryonic stem cells.

To improve the scDamID method and make it compatible with simultaneous mRNA measurement in single cells, we optimized several shortcomings of the previously developed protocol^2^. The improvements include (1) the requirement of one, rather than two ligation events to amplify fragmented gDNA molecules, (2) switching from PCR to linear amplification through *in vitro* transcription, (3) inclusion of unique molecule identifiers (UMI) for both gDNA- and mRNA-derived reads, and (4) the use of liquid-handling robots that result in rapid and higher processing throughputs of thousands of single cells per day together with reduced reaction volumes, and a more consistent sample quality. As described previously^2^, KBM7 cells (a near haploid myeloid leukemia cell line, except for chr8 and parts of chr15) expressing either untethered Dam or a Dam-LmnB1 fusion protein and the 2-colour Fucci reporter system^18^ are sorted by FACS at the G1/S cell cycle transition 15 hours post-induction of Dam with Shield1^2^. After single cells are sorted into 384-well plates, poly-adenylated mRNA is reverse transcribed using primers that contain a T7 promoter, P5 Illumina adapter, a random UMI sequence, and mRNA- and cell-specific barcodes in the overhang, as described previously for the CEL-Seq protocol^19-20^ (Fig. 1a). Second strand synthesis is then performed to generate double-stranded cDNA. Next, the reaction mixture, containing tagged cDNA molecules and gDNA, is digested with the restriction enzyme DpnI. DpnI recognizes adenine residues that are methylated by Dam in a GATC context and creates blunt double-stranded cuts in gDNA. Double-stranded adapters are then ligated to digested gDNA molecules (Fig. 1a). Similar in design to the RT primers, the double-stranded adapters contain a T7 promoter, P5 Illumina adapter, UMI, and gDNA- and cell-specific barcodes. Single cells are then pooled, and cDNA and ligated gDNA molecules, both containing T7 promoter sequences, are simultaneously amplified by *in vitro* transcription. The amplified RNA molecules are then used to prepare Illumina libraries, as described previously^20^ (Fig. 1a). Thus, this new method enables genome-wide quantification of protein-DNA interactions and mRNA from the same cell without requiring physical separation steps, thereby minimizing losses and making it easily adaptable to automated liquid handlers that can process thousands of single-cells per day in a high-throughput format.

**Figure 1.**
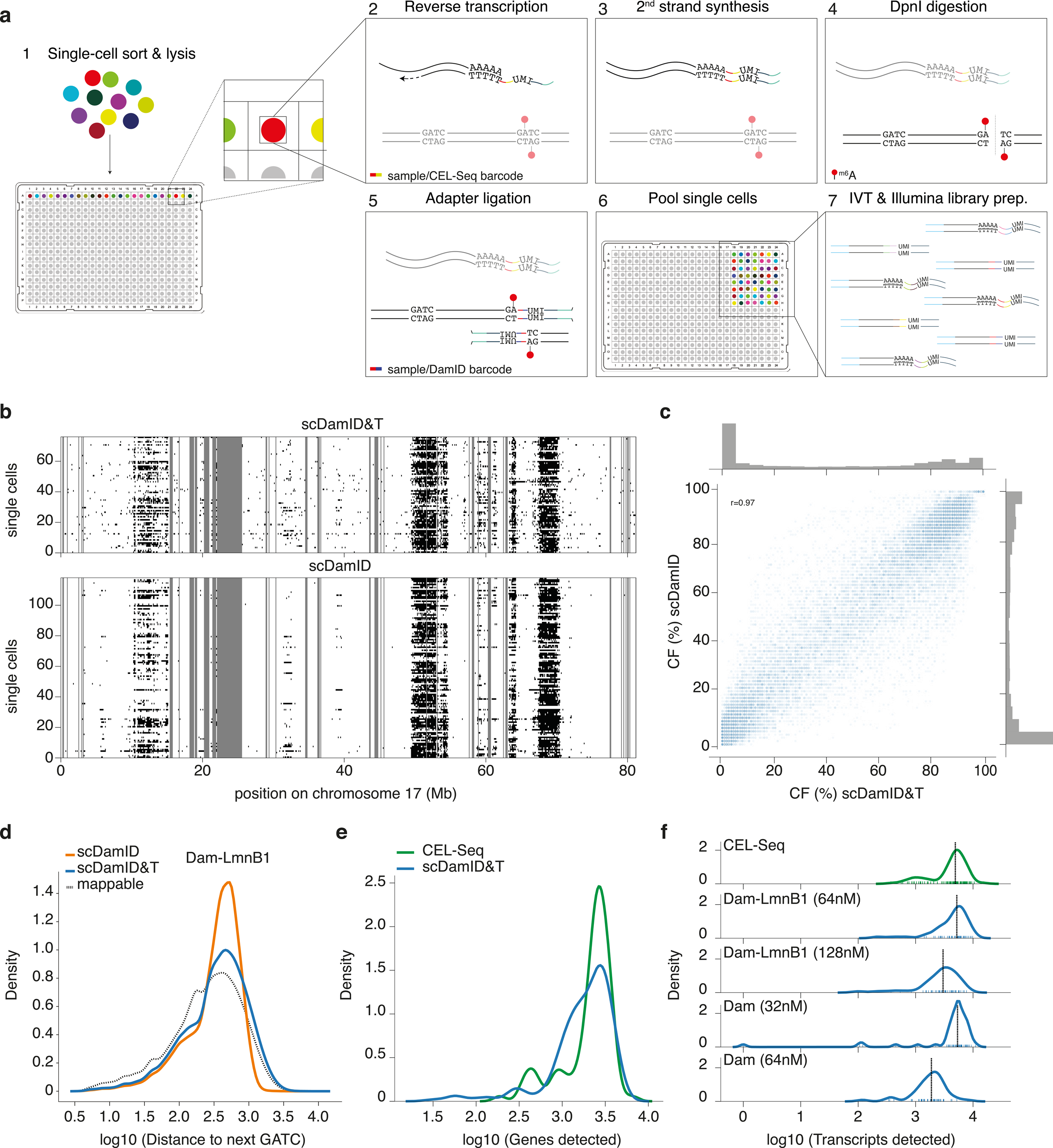
Quantitative comparison of scDamID, CEL-Seq and scDamID&T applied to KBM7 cells. **a)** Schematic representation of the scDamID&T method. **b)** Binary representation of OE values of Dam-LmnB1 signal measured with scDamID&T and scDamID^2^ in single cells on chromosome 17. Unmappable regions are marked in grey. **c)** Comparison of CFs for scDamID (y-axis) and scDamID&T (x-axis). CF distributions are depicted in the margins. Pearson’s r = 0.97. **d)** Distribution of inter-GATC distances of mappable GATC fragments genome-wide (dotted line), and inter-GATC distances of GATCs observed with scDamID (orange line) and scDamID&T (blue lines) for Dam-LmnB1. **e)** Distributions of the number of unique genes detected using CEL-Seq^2^ (green line) and scDamID&T (blue line). **f)** Distribution of the number of unique transcripts detected by CEL-Seq data^2^ (green line) and scDamID&T (blue line) for Dam and Dam-LmnB1, and for different DamID adapter concentrations.

To determine the efficiency of the combined method, we benchmarked scDamID&T to previous data in KBM7 cells; a clonal line for which single-cell genome-NL interaction maps (scDamID) and single-cell transcriptomes are already available^2^. We successfully detected reads corresponding to both DamID and mRNA. We detected a median of 60,348 unique DamID reads per cell, identifying all major LADs, as previously reported from bulk and single-cell sequencing^2^. As illustrated for chromosome 17, observed over expected (OE) scores^2^ calculated based on the combined method not only detected all LADs but also captured the cell-to-cell heterogeneity in genome-NL interactions as observed previously (Fig. 1b and Supplementary Fig. 1a). This is further illustrated by the high concordance (Pearson *r* = 0.97) in the contact frequencies (CFs), the percentage of cells, which at a given position in the genome are in contact with the NL (Fig. 1c). Altogether this shows that scDamID&T can successfully capture the dynamics of genome-NL interactions in single cells. A crucial improvement in the scDamID&T method is that the cell- and nucleic acid-specific barcoding enables single cells to be pooled prior to amplification and library preparation, as opposed to the individual cell library preparation and sample selection in scDamID. This significantly contributes to increased throughput and cost reduction. Although single cells are pooled in scDamID&T prior to amplification without selection for cells with the highest signal, the complexity of the single-cell libraries, quantified as the number of unique reads per read sequenced in a cell, is comparable between both methods (Supplementary Fig. 1b). Further, the loss of reads with incorrect adapter sequences is substantially reduced in the new method (Supplementary Fig. 1c). The previously developed scDamID is biased against detection of GATC sites that were separated by over 1 kb in the genome; a drawback that is overcome by a single ligation event in scDamID&T which captured the genome-wide distribution of GATC sites more faithfully (Fig. 1d and Supplementary Fig. 1d).

Next, we benchmarked the transcriptomic measurements from scDamID&T to previously obtained single-cell CEL-Seq data for KBM7 cells^2^. Both methods detected the expression of comparable number of genes (Median: CEL-Seq = 2509, scDamID&T = 2052) (Fig. 1e), and the number of unique transcripts detected per cell was similar for both methods (Median: CEL-Seq = 4920, scDamID&T = 3743) (Supplementary Fig. 2a). The efficiency of mRNA detection appears to reduce with higher DamID double-stranded adapter concentrations; we find that the quality of the transcriptome libraries can be further increased by lowering the double-stranded adapter concentrations, without compromising the quality of the DamID libraries (Fig. 1f and Supplementary Fig. 2b). Hierarchical clustering of the single-cell transcriptomes showed that samples from both methods cluster together (Supplementary Fig. 2c), emphasizing the concordance between the transcriptomes captured by both techniques.

To verify scDamID&T in an independent cell line, we also established the system in hybrid (129/Sv:Cast/EiJ) mouse embryonic stem (mES) cells^21^ where DamID expression is controlled via the auxin-AID degron system^22^ (Supplementary Fig. 3a). The quality of the scDamID&T libraries in mES cells expressing Dam or Dam-LmnB1 is comparable to KBM7 cells except that the single-cell Dam-LmnB1 data is of lower complexity (Supplementary Fig. 3b). The reduction in DamID complexity is likely a reflection of the shorter induction time of Dam-LmnB1 in mES cells and difference in cell cycle characteristics. Nevertheless, measurements with scDamID&T from these samples show strong DamID signals in previously reported^23^ bulk LAD domains (Supplementary Fig. 3c).

Extrapolating the technology that we developed for the detection of genome-NL interactions and mRNA from the same cell, we hypothesized that KBM7 cells expressing untethered Dam could be used to quantify both DNA accessibility and the transcriptome on a genome-wide scale from single cells. To explore the possibility of using Dam as a DNA accessibility marker, we first quantified the levels of Dam GATC methylation of averaged single-cell profiles around transcription start sites (TSS) of actively transcribed genes and observed a sharp peak at these sites (Fig. 2a). As a control, we also performed these single-cell experiments using the non-methylation sensitive restriction enzyme AluI. We did not observe signatures of accessibility around TSS of actively expressed genes (Fig. 2b), indicating that the observed Dam accessibility patterns are the result of *in vivo* Dam methylation at accessible regions of the genome, and not a consequence of restriction enzyme accessibility. Similar to active TSSs, we also observe strong Dam enrichment at active enhancers (Fig. 2c).

**Figure 2.**
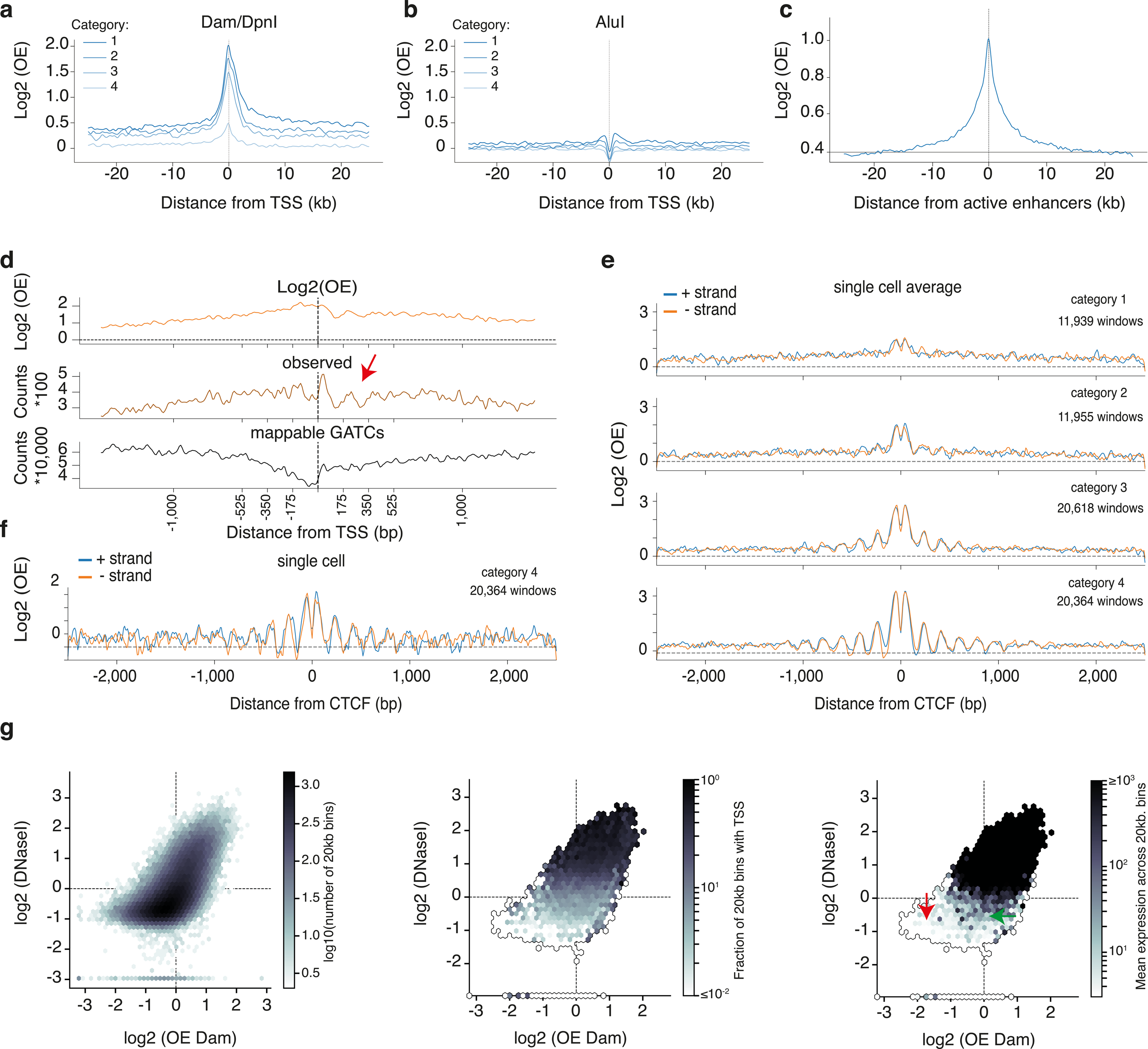
Untethered Dam marks accessible chromatin in single cells. **a)** Transcription start site (TSS) alignment of the single-cell average (n=96 cells) Dam signal stratified by gene expression into four categories of expression levels (category 1 most active or highly expressed; category 4 least active or not expressed). **b)** TSS alignment as for (a), showing the density of AluI-derived genomic fragments. **c)** Alignment plot of the single-cell average (n=96 cells) Dam signal at active enhancers. **d)** TSS alignment of the single-cell average (n=96 cells) Dam signal for active genes at 10bp resolution for OE values (orange), observed reads (brown) and density of mappable GATCs (black). The red arrow highlights an example of periodicity in the DNA accessibility signal. **e)** Single-cell average (n=96 cells) Dam signal alignment at CTCF sites, stratified in four regimes of increasing CTCF binding activity (see computational methods for details on stratification). **f)** Example of Dam signal at CTCF sites for a single cell with the highest CTCF binding activity. **g)** Scatter plot of bulk DNaseI (y-axis) and single-cell average Dam data (x-axis). The left panel displays the density of 20kb bins as a function of DNaseI (y-axis) and Dam (x-axis) signal. The middle panel displays the density of 20kb bins with at least a single TSS. The right panel depicts the mean expression for all genes in all 20kb regions for each point in the plot. Note that for baseline DNaseI signal (red arrow), genes that are expressed at low levels display elevated Dam signal (green arrow).

Nucleosomes are known to be regularly spaced on active TSS^24,25^ and CTCF sites, and this can be observed in DNA accessibility data pooled across 96 single cells obtained using scDamID&T (Fig. 2d and 2e and Supplementary Fig. 4a). The observed periodicity of 178bp is in general agreement with the reported spacing of nucleosomes in human cells^25^ (Supplementary Fig. 4b). Remarkably, these nucleosome positioning profiles are also apparent in data from single cells (Fig. 2f), indicating that Dam can serve to determine nucleosome positioning *in vivo* in single cells. This feature could be especially powerful when scDamID&T is combined with single-cell CRISPR/Cas9 to screen for factors involved in nucleosome positioning^26^. When comparing Dam-mediated DNA accessibility data to bulk DNaseI-seq data, we find that the dynamic range of Dam-mediated DNA accessibility is larger; for a substantial fraction of the genome only baseline levels of DNaseI are detected, while Dam indicates intermediate levels of accessibility (Fig. 2g). Further analyses showed that these regions are typified by genes with low expression, indicating that Dam is more sensitive than DNaseI and allows discrimination between inactive and lowly transcribed genes. This feature may be attributed to the advantage of Dam detecting both active promoters (H3K4me3) and gene bodies (H3K36me3) (Supplementary Fig. 4c) and the *in vivo* accumulation of Dam signal over time.

As scDamID&T enables simultaneous quantification of protein-DNA interactions and mRNA from the same cell, we next investigated how variations in genome-NL association directly influence gene expression. Further, as dissociation of genomic loci from the NL has been shown to result in an increase in active histone modifications for some of those loci ^17^, we hypothesized that the propensity of a region in the genome to associate with the NL could result in differentially regulated gene expression. To test this hypothesis, we first quantified heterogeneity in genome-NL associations for each 500 kb region using CFs^2^. While single-cell samples generally show a large degree of concordance, certain regions are found in contact with the NL in only a small fraction of cells (“low CF”). We found that gene expression in that small fraction of cells that exhibit NL contact is generally lower compared to cells that do not show NL contact (for example genomic region 839, Fig. 3a). In contrast, for regions with intermediate CF (for example genomic region 317, Fig. 3a), gene expression was independent of NL-positioning (Fig. 3a “middle CF”). Performing this analysis on a genome-wide scale and stratifying bins by their CF values, we found a significant decrease of gene expression upon NL association in regions with low CF values (Fig. 3b), whereas genomic regions with CF values greater than 20% appear to be insensitive to NL association. Interestingly, the impact on gene expression does not seem to vary with the (mean) gene expression levels (Supplementary Fig. 5a). Taken together, these results suggest that the CF of a region biases the sensitivity of gene expression to NL positioning. To our knowledge, this is the first report to show that heterogeneity in spatial positioning of the genome directly impacts gene expression in single cells. Finally, this differential sensitivity in transcriptional output of genomic regions upon NL association may explain the varied outcomes of three previous studies showing that artificial targeting of genomic regions to the NL resulted in reduced, mixed or unchanged expression levels of the genes^27-29^.

**Figure 3.**
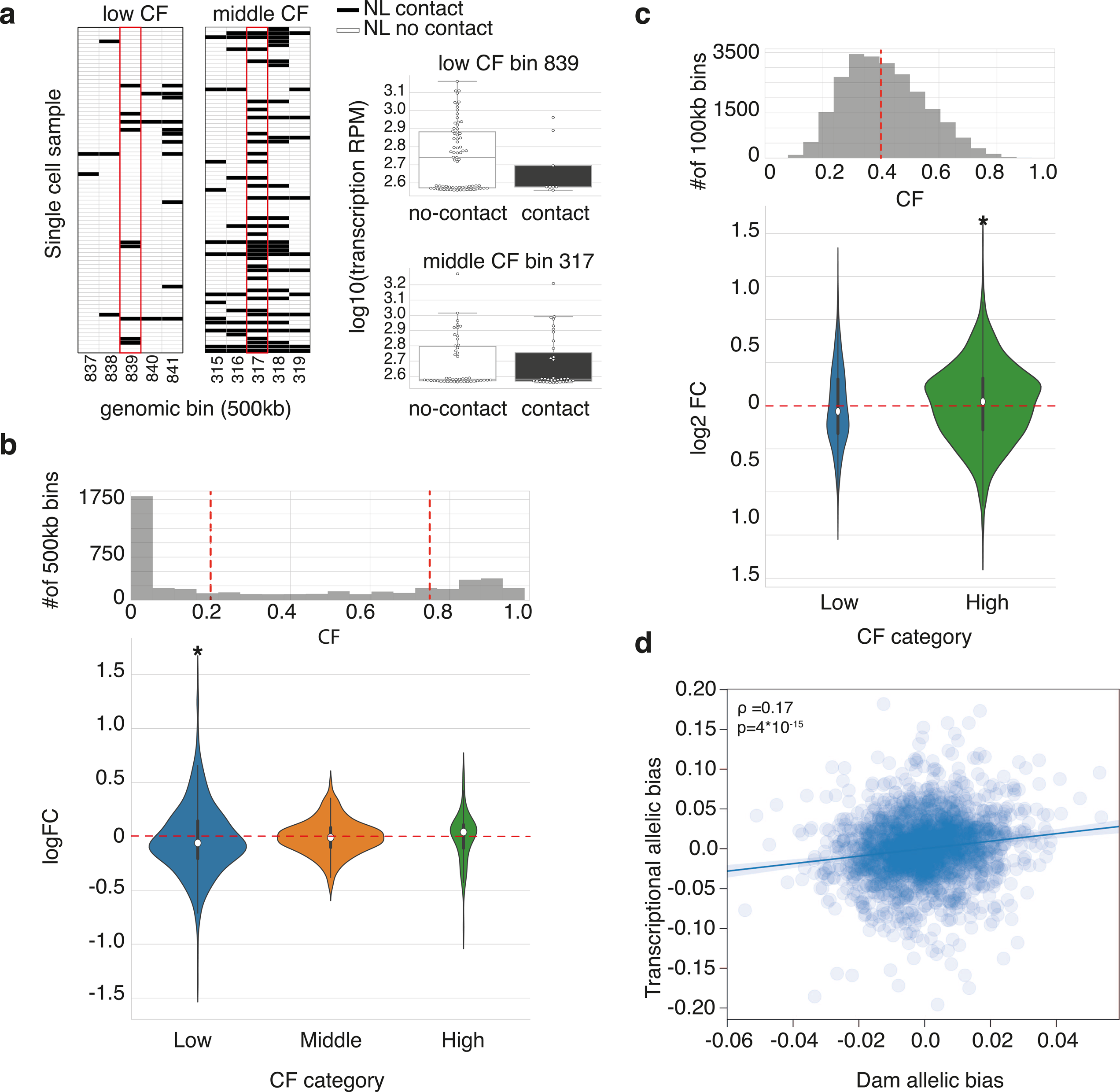
Parallel transcriptomic and DamID measurements link transcriptional dependencies with heterogeneity in DamID contacts. **a)** Examples of regions with low (left) and intermediate (right) CFs. The black filled boxes indicate single-cell 500kb NL contacts (OE value > 1); white boxes indicate no NL contact (OE value < 1). Boxplots in the right panels display gene expression levels in these bins, stratified by NL contacts. For the low CF bin, note the increased expression levels in cells with no NL contacts. Bin 839 corresponds to genomic region chr2:170000000-170500000. Bin 317 corresponds to genomic region chr1:158500000-159000000. **b)** Top panel: distribution of CF values across the genome for Dam-LmnB1 data in KBM7 cells. Red lines indicate the segmentation of the genomic regions in low, intermediate and high CF bins. Bottom panel: distributions of log2 fold-change (FC) in gene expression between cells exhibiting contact *vs.* cells not exhibiting contact. *=p<0.05, two-sided t-test. **c)** Analysis as in b, for untethered Dam in KBM7 cells. *=p<0.05, two-sided t-test. **d)** Scatter plot of the measured mES cell allelic bias (129/Sv vs. Cast/EiJ) in transcription (y-axis) *vs.* the allelic bias in chromatin accessibility (x-axis), measured in 100kb bins. Chromosomes 5, 8 and 12, as well as the sex chromosomes were excluded from this analysis.

Next, we applied this analysis to explore how variability in DNA accessibility relates to heterogeneity in gene expression in KBM7 cells. We found that for regions that were in contact with Dam in a large fraction of the cells (CF > 40%), expression was significantly higher in cells showing Dam contact (Fig. 3c and Supplementary Fig. 5b). These results suggest that gene expression heterogeneity between single cells is more sensitive to variability in DNA accessibility within open chromatin regions. Consistent with the results of KBM7 cells, we also observed the same relationship in the hybrid mES cells, suggesting that the observed relationship between DNA accessibility and gene expression is generalizable to other mammalian systems (Supplementary Figs. 5c and 5d)

To expand upon the analysis presented above, we investigated how DNA accessibility tunes gene expression at an allelic resolution. For this, we used a hybrid mES cell line of 129/Sv:Cast/EiJ genotype^21-30^ which is known to harbor a duplication of Cast/EiJ chromosome 12. In order to carefully karyotype this cell line prior to application of scDamID&T, we modified our technique to detect copy number variations in single cells, by using the Dam-methylation insensitive restriction enzyme AluI instead of DpnI. This demonstrates that scDamID&T can also be easily extended to quantify the genome and transcriptome from the same cell, using minor modifications to the protocol presented above^13,14^. The AluI data showed that the hybrid mES cell line harbors a systematic duplication of the Cast/EiJ chromosome 12 in most but not all single cells (Supplementary Fig. 6a). When we performed scDamID&T using untethered Dam to measure single-cell DNA accessibility profiles we also detected increased Dam contacts for the Cast/EiJ chromosome 12, and a chromosome-wide mRNA bias towards Cast/EiJ transcripts (Supplementary Fig. 6b and 6c). Surprisingly, we also detected a small fraction of cells that displayed increased DNA accessibility for the 129/Sv allele over the Cast/EiJ allele for chromosome 12, and a corresponding increase in 129/Sv derived transcripts for one cell (Supplementary Figure 6c). After excluding the confounding effects of CNVs on chromosome 12 as well as chromosomes 5 and 8 in this hybrid mES cell line, we observed a significant positive correlation between allele-specific DNA accessibility and gene expression (Fig. 3d). Taken together, these results demonstrate that scDamID&T can also be used to directly quantify the allele-specific relationship between DNA accessibility and the transcriptome (Supplementary Figs. 6a-c).

Finally, we sought to test scDamID&T as an *in silico* cell sorting strategy to distinguish and group cell types based on the transcriptomes and thereafter, uncover the underlying cell-type specific gene-regulatory landscape by DamID. Such a strategy to obtain cell-type specific protein-DNA interaction maps is particularly attractive for complex tissues and tumors with unknown cellular constitution, or for certain cell types that cannot be isolated with sufficient purity due to a lack of discriminating surface markers or a lack of high quality antibodies.

To demonstrate that our new technology can be used as an *in silico* cell sorting technique that enables generation of cell-type specific DNA accessibility profiles, we performed a proof-of-principle experiment where mES cells cultured under 2i or serum conditions were sorted and quantified using scDamID&T. Single-cell transcriptomes obtained using scDamID&T could be used to readily separate the population into two distinct clusters, corresponding to 2i and serum grown cells (Fig. 4a). Expression analysis showed signature genes differentially expressed between the two conditions (Supplementary Fig. 7a). DNA accessibility profiles generated from the two *in silico* transcriptome clusters showed differential accessibility patterns on a genome-wide scale. For example, DNA accessibility tracks along Peg10, a gene strongly upregulated under serum conditions, showed increased accessibility at the TSS and along the length of the gene (Fig. 4b). Interestingly, the increased accessibility in the serum condition extends beyond the Peg10 gene locus, encompassing the entire length of a large topologically associated domain (TAD). Indeed, the overall expression of neighboring genes within this TAD is higher in serum conditions (Fig. 4b). Generalizing this to all differentially expressed genes, we found that upregulation of gene expression in 2i or serum conditions correlated with increased DNA accessibility over the entire gene body (Figs. 4c and 4d and Supplementary Fig. 7b). Similarly, we observed that differentially upregulated genes in each condition showed an increase in DNA accessibility at the TSS for those genes (Fig. 4d). Thus, these results demonstrate that scDamID&T can be used to effectively generate cell-type specific DNA accessibility profiles. Finally, we found that upregulated gene expression also correlated with increased accessibility at the single-cell level, highlighting that scDamID&T can be used to study changes in cellular identities in direct relationship with the accompanying gene-regulatory mechanisms that shape cell type-specific gene expression programs (Fig. 4e).

**Figure 4.**
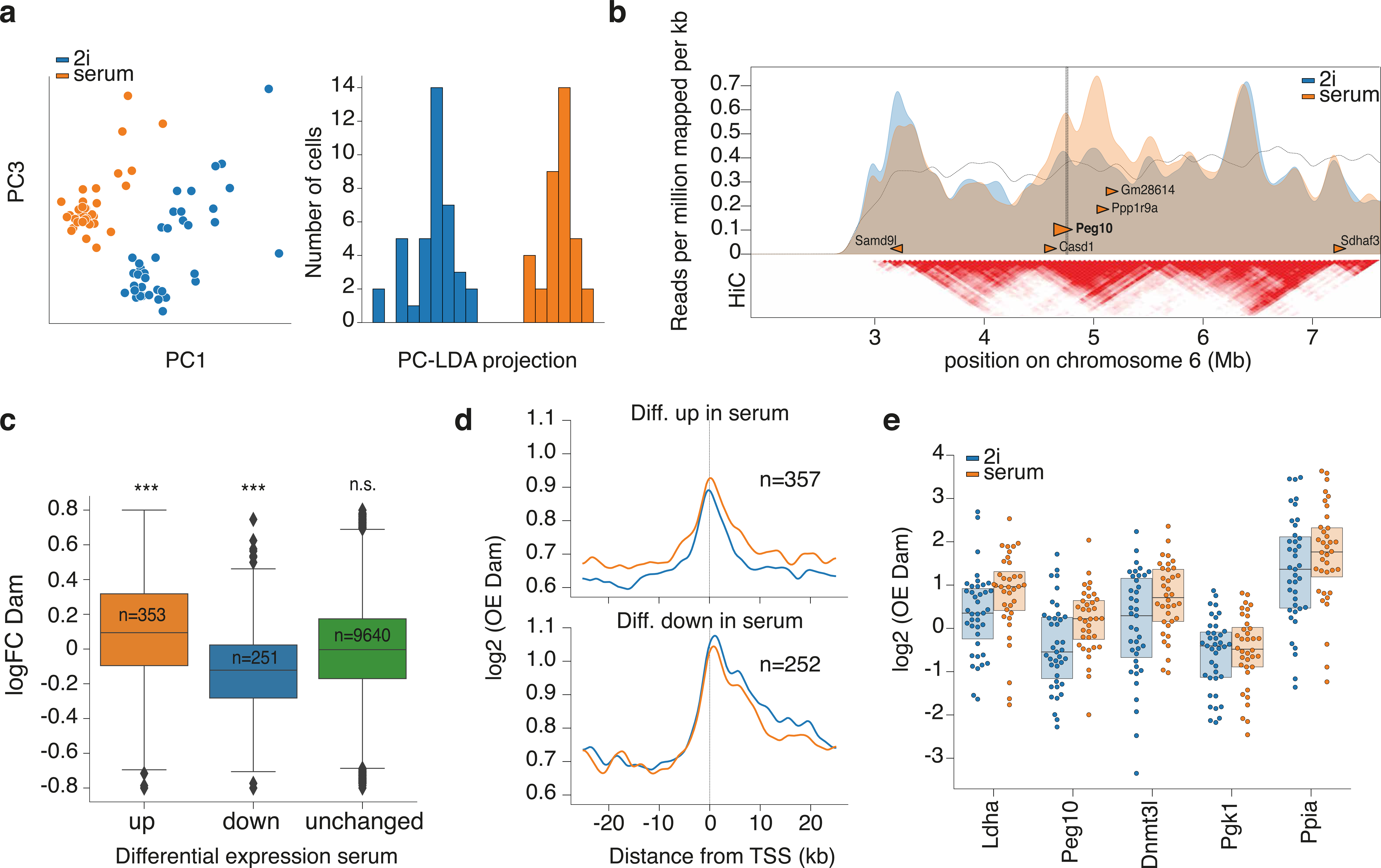
scDamID&T enables *in silico* cell sorting and reconstruction of corresponding cell type specific gene regulatory landscapes. **a)** Principle component (left) and principal components-linear discriminant (right) analysis on Dam expressing mES cells cultured in 2i (blue) or serum conditions (orange). **b)** DNA accessibility profiles in 2i and serum conditions. Arrowheads indicate genes with log2FC of ≥1.25 in serum condition. Arrowheads with black outline were found to be significantly differentially expressed (with FDR < 5%). The lower panel shows HiC data obtained from mESCs^32^ displayed with the 3D genome browser {DOI:<u>10.1101/112268</u>}. **c)** log2 FC in DNA accessibility between serum and 2i conditions for genes that are differentially up (orange), down (blue) or unaffected (green) in serum conditions compared to 2i. **d)** DNA accessibility at TSSs of differentially up-(top panel) or down-regulated (bottom panel) genes in serum (orange line) conditions compared to 2i (blue line). **e)** DNA accessibility for the top 5 induced genes in serum compared to 2i condition in single cells (cells are represented by dots).

In summary, we have developed a new technology to simultaneously quantify genome-NL interactions (Dam-LmnB1), DNA accessibility (Dam) or genome CNVs (AluI) with the transcriptome from the same cell. scDamID&T enables dissection of the relationship between the direct impact of spatial genome organization and chromatin accessibility on gene expression. Further, it can be applied to sort cell types *in silico* and obtain their associated gene-regulatory landscapes. Excitingly, in the future, scDamID&T can be employed to obtain combined single-cell quantifications of many distinct nuclear regulatory mechanisms via the coupling of Dam to transcription factors, various constituents of different chromatin types (for example, Polycomb-group proteins and HP1) or the DNA replication or DNA damage machineries^31^. Applied to dynamic biological processes, this technique should prove especially powerful to dissect the order and sequence of epigenetic changes that are necessary for the acquisition of different cell fates in heterogeneous tissues and differentiation systems.

## Acknowledgements

We would like to thank the members of the JK and AvO labs for their comments on the manuscript. We would also like to thank Mauro Muraro and Lennart Kester for valuable input setting up this technique. This work was supported by an European Research Council Starting grant (ERC-STG 678423-EpiID), Advanced grant (ERC-AdG 742225-IntScOmics) and a Netherlandse organisatie voor Wetenschappelijk Onderwijs (NWO) open grant (824.15.019) and TOP award (NWO-CW 714.016.001).

## Competing interests statement

The authors declare that they have no competing financial interests.

Correspondence and requests for materials should be addressed to S.S.D. or J.K.

## Data availability

The sequencing DamID data from this study are available from the Gene Expression Omnibus, accession number GSE108639 (https://www.ncbi.nlm.nih.gov/geo/query/acc.cgi?acc=GSE108639). The data can be accessed with the use of the token: ytsvcsiqhzoppux.

---

**Supplementary Figure 1 | Quantitative comparison between scDamID and ScDamID&T**

**a)** Comparison between the binarized single cell (horizontal tracks) contact frequency maps for scDamID (top panel 118 cells) and scDamID&T (bottom panel 93 cells) **b)** Comparison of sample complexities with scDamID (orange) and scDamID&T (blue) depicted by unique reads (y-axis) with increasing sequencing depth (x-axis) in single-cell samples. **c)** Overview of losses during processing of raw sequencing data in scDamID (orange bars) and scDamID&T (blue bars). The raw reads are first filtered on the correct adapter structure, then aligned to the human genome, where reads not yielding a unique alignment are filtered out, as well as reads not aligning immediately adjacent to GATCs. Finally, duplicate reads are removed, on account of the haploid nature of the KBM7 cell-line. **d)** Distribution of inter-GATC distances of mappable GATC fragments genome-wide (dotted line), and inter-GATC distances of GATCs observed with scDamID (orange line) and scDamID&T (blue lines) for Dam.

**Supplementary Figure 2 | Quantitative comparison between CEL-Seq and scDamID&T**

**a)** Distributions of the number of unique transcripts detected using CEL-Seq^2^ (green line) and scDamID&T (blue line). **b)** Overview of losses during processing of transcriptomic data obtained with CEL-Seq (green bars) or scDamID&T (blue bars). The raw reads are aligned to the human genome, reads that do not yield unique alignments are filtered, as well as reads that do not match exons. Finally, duplicate reads are removed based on the UMIs. **c)** Hierarchical clustering of the transcriptomes obtained with CEL-Seq (green) and scDamID&T (blue).

**Supplementary Figure 3 | ScDamID&T in hybrid mES cells**

**a)** Auxin mediated control of AID-Dam and AID-Dam-LmnB1 cell lines. DamID PCR products of cells 24- and 48hours after auxin washout (top panel). Time course and quantitative PCR analysis of auxin induction for a locus within a LAD, 0-, 8-, 10-, 12- and 24 hours after auxin washout (bottom panel). Quantification of the ^m6^A levels as described for the DpnII assay^17^. **b)** Overview of losses during data processing as in Supplementary Figure 2a for the scDamID&T libraries obtained in mES cells. **c)** mES Dam-LmnB1 OE values projected on the upstream (top panel) and downstream (bottom panel) of LAD-boundaries defined previously^23^.

**Supplementary Figure 4 | Untethered Dam enzyme marks accessible chromatin in single cells**

**a)** TSS alignment of the single-cell average (n=96 cells) Dam signal for inactive genes at 10bp resolution for OE values (orange), observed reads (brown) and mappable GATCs (black). **b)** 10bp resolution frequency spectrum of single-cell average (n=96 cells) Dam-signal stratified in four regimes of increasing CTCF binding activities. Note the peak signal for the CTCF sites with the highest binding activities corresponds to 178bp (red arrow). **c)** Distribution of 20kb bins as function of bulk H3K4me3 (y-axis, left panel) or bulk H3K36me3 (y-axis, right panel) and single-cell average Dam data (x-axis). Increasing grey-level intensity represents increasing 20kb bin density.

**Supplementary Figure 5 | Single-cell associations between transcription and Dam or Dam-LmnB1 contacts**

**a)** log2 FCs in expression levels (y-axis) between Dam-LmnB1 contact (OE > 1) and no contact (OE < 1) samples, measured in 500kb bins, versus log-scaled expression levels (x-axis). Note that negative log2 FCs indicate higher expression in the “no NL-contact” samples compared to “NL-contact” samples. The dotted line indicates a locally-weighted regression (“lowess”). **b)** log2 FCs in expression levels (y-axis) calculated between contact and no contact samples in KBM7 cells expressing untethered Dam, as in **a**. Note that positive log2 FCs indicate higher expression in the “Dam contact” samples compared to the “no Dam contact” samples. **c)** Violin plot for the log2 FC expression levels between contact and no-contact samples obtained with Dam-expressing hybrid mES cells, as Fig. 3b and Fig. 3c. *=p<0.05, two-sided t-test. **d)** Same as for **b**, but in Dam expressing hybrid mES cells.

**Supplementary Figure 6 | Allelic associations between single-cell transcription and Dam contacts**

**a)** AluI signal obtained from 74 129/Sv:Cast/Eij mES cells. Each row represents a single cell; each column a 100kb bin along the genome. The checkered black box indicates the duplication of the Cast/EiJ chromosome 12. The track below the plot shows allelic bias for the maternal 129/Sv allele in purple and the paternal Cast/EiJ allele in green, as determined using partial least squares regression. **b)** Plot as in A, showing DamID signals obtained from 67 129/Sv:Cast/EiJ mES cells. **c)** Allelic bias in transcription (y-axis) in relationship to the allelic bias in Dam signal (x-axis) for chromosome 12. One single cell (named #12) exhibits about 2-fold lower Dam signal and transcriptional output from the Cast/EiJ allele (right panel), while exhibiting a 2-fold increase in Dam and transcriptional signals originating from the 129/Sv allele (left panel).

**Supplementary Figure 7 | *In silico* sorting of cell identities and corresponding regulatory landscapes with scDamID&T**

**a)** log2-transformed expression values for the top five differentially up-regulated genes in 2i (left) and serum (right) conditions. The horizontal line for Gpx2 in serum conditions indicates no expression. **b)** Density plot of genes relating the log2 FC in Dam accessibility (x-axis) to log2 FC in gene expression (y-axis), showing only genes that were found to be differentially expressed between 2i and serum conditions (FDR < 5%).

**Supplementary table 1 | scDamID double-stranded adapters**

**Supplementary table 2 | CEL-Seq2 primers**

**Supplementary table 3 | Statistical details per figure**

## Methods

Cell culture. Haploid KBM7 cells were cultured in suspension in IMDM (Gibco) supplemented with 10% FBS and 1% Pen/Strep. The same Shield1-inducible Dam-LmnB1 and Dam-only stable clonal KBM7 cell lines were used as in ^1^. Cells were split every 3 days. F1 hybrid 129/Sv:Cast/Eij mouse embryonic stem cells (mESCs)^2^ were cultured on primary mouse embryonic fibroblasts (mEFs), in ES cell culture media; G-MEM (Gibco) supplemented with 10% FBS, 1% Pen/Strep, 1x GlutaMAX (Gibco), 1x non-essential amino acids (Gibco), 1x sodium pyruvate (Gibco), 143 µM β-mercaptoethanol and 1:1000 hLIF (in-house production). Cells were split every 3 days. Expression of constructs was suppressed by addition of 0.5 µM and indole-3-acetic acid (IAA; Sigma, I5148). 2i F1 hybrid 129/Sv:Cast/Eij mESCs cells were cultured for 2 weeks on primary mEFs in 2i ES cell culture media; 48% DMEM/F12 (Gibco) and 48% Neurobasal (Gibco), supplemented with 1x N2 (Gibco), 1x B27 supplement (Gibco), 1x non-essential amino acids, 1% Pen/Strep, 143 uM β-mercaptoethanol, 0.5% BSA, 1 µM PD0325901 (Axon Medchem, 1408), 3 µM CHIR99021 (Axon Medchem, 1386) and 20 ng/mL hLIF (in-house production). Cells were split every 3 days. Expression of constructs was suppressed by addition of 0.5 µM IAA.

### Generating cell lines

Stable clonal Dam and Dam-LmnB1 F1 hybrid mESC lines were created by co-transfection of the EF1alpha-Tir1-neo and hPGK-AID-Dam-mLmnb1 or hPGK-AID-Dam plasmids in a ratio of 1:5. Cells were trypsinized and 0.5 × 10^6^ cells were plated directly with Effectene transfection mixture (Qiagen, 301427) on 0.1% gelatin (in-house production) in 60% BRL-conditioned medium. The transfection was according to the kit protocol. Cells were selected for 10 days with 250 µg/mL G418 and selection of the clones was based on methylation levels, determined by DpnII-qPCR assays as previously described ^3^ To reduce the background methylation levels in the presence of 1.0 mM IAA (Sigma, I5148), we transduced the selected clones of both AID-Dam-LmnB1 and Dam-only with extra hPGK-Tir1-puro followed by selection with 0.8 µg/mL puromycin. Positive clones were screened for IAA induction in the presence and absence of IAA by DpnII-qPCR assays and DamID PCR products.

### DamID induction

Expression of Dam-LmnB1 or Dam-only constructs was induced in the KBM7 cells with 0.5 nM Shield1 (Glixx laboratories, 02939) 15 hours prior to harvesting as described previously ^1^. Expression of Dam-LmnB1 or Dam-only constructs was induced in the F1 mESCs by IAA washout 12 hours prior to harvesting. Based on the growth curve of cells counted at time points 0, 12, 24, 30, 36, 42, 48, 54, 60, 72 and 84 after plating, the generation time of both the Dam-LmnB1 and Dam-only cell lines was estimated at ∼12 hours (data not shown). Considering that 55% of the cells are in G1 and early S, the estimated time these cells reside in G1 and early S is 6,75 hours.

### Cell harvesting and sorting

KBM7 cells were harvested in PBS (in-house production), stained with 0.5 μg/mL DAPI for live/dead selection. Small haploid Single cells were sorted based on forward and side-scatter properties (30% of total population) and selected for double positive FUCCI profile as described before ^1^ F1 mES cells were collected in plain or 2i ES cell culture media, stained with 30 μg/mL Hoechst 34580 for 45 minutes at 37°C. mES cell singlets were sorted based on forward and side-scatter properties, and in mid-S phase of the cell cycle based on DNA content histogram. One cell per well was sorted into 384-well plates (Biorad, HSP3801) using the BD FACSJazz cell sorter. Wells contained 4 µL mineral oil (Sigma) and 100 nL of 15 ng/µL unique CELseq primer.

### scDamID&T

Robotic preparation: 4 µL mineral oil was dispensed manually into each well of a 384-well plate using a multichannel pipet. 100 nL of unique CEL-seq primer was dispensed per well using the mosquito HTS robot (TTP Labtech). The NanodropII robot (BioNex) was used for all subsequent dispensing steps at 12 p.s.i. pressure. After sorting, 100 nL lysis mix was added (0.8 U RNase inhibitor (Clontech, 2313A), 0.07% Igepal, 1mM dNTPs, 1:500000 ERCC RNA spike-in mix (Ambion, 4456740)). Each single cell was lysed at 65°C for 5 min and 150 nL reverse transcription mix was added (1x First Strand Buffer (Invitrogen, 18064-014), 10 mM DTT (Invitrogen, 18064-014), 2 U RNaseOUT Recombinant Ribonuclease Inhibitor (Invitrogen, 10777019), 10 U SuperscriptII (Invitrogen, 18064014)) and the plate was incubated at 42°C for 1 h, 4°C for 5 min and 70°C for 10 min. Next, 1.92 µL of second strand synthesis mix was added (1x second strand buffer (Invitrogen, 10812014), 192 µM dNTPs, 0.006 U *E. coli* DNA ligase (Invitrogen, 18052019), 0.013 U RNAseH (Invitrogen, 18021071)) and the plate was incubated at 16°C for 2 h. 500 nL of protease mix was added (1x NEB CutSmart buffer, 1.21 mg/mL ProteinaseK (Roche, 000000003115836001)) and the plate was incubated at 50°C for 10 hr and 80°C for 20 min. Next, 230 nL DpnI mix was added (1x NEB CutSmart buffer, 0.2 U NEB DpnI) and the plate was incubated at 37°C for 4 hr and 80°C for 20 min. Finally, 50 nL of DamID2 adapters were dispensed (final concentrations varied between 2 and 128 nM), together 450 nL of ligation mix (1x T4 Ligase buffer (Roche, 10799009001), 0.14 U T4 Ligase (Roche, 10799009001)) and the plate was incubated at 16°C for 12 hr and 65°C for 10 min. Contents of all wells with different primers and adapters was pooled and incubated with 0.8x 1:4 diluted magnetic beads (CleanNA, CPCR-0050) for 10 min, washed twice with 80% ethanol and resuspended in 7 µL nuclease-free water before *in vitro* transcription at 37°C for 14 hr using the MEGAScript T7 kit (Invitrogen, AM1334). Library preparation was done as described in the CEL-seq protocol with minor adjustments ^4^. Amplified RNA (aRNA) was cleaned and size-selected by incubating with 0.8x magnetic beads (CleanNA, CPCR-0050) for 10 min, washed twice with 80% ethanol and resuspended in 22µL nuclease-free water, and fragmented at 94°C for 2 min in 0.2x fragmentation buffer (200 mM Tris-acetate, pH 8.1, 500 mM KOAc, 150 mM MgOAc). Fragmentation was stopped by addition of 0.1x fragmentation STOP buffer (0.5 M EDTA pH8) and quenched on ice. Fragmented aRNA was incubated with 0.8x magnetic beads (CleanNA, CPCR-0050) for 10 min, washed twice with 80% ethanol and resuspended in 12 µL nuclease-free water. Thereafter, library preparation was done as previously described ^4^ using 5 µL of aRNA and PCR cycles varied between 8 and 10. Libraries were run on the Illumina NextSeq platform with high output 75bp paired-end sequencing.

### DamID adapters

The adapter was designed (5’ to 3’) with a 4 nt fork, a T7 promoter, the 5’ Illumina adapter (as used in the Illumina small RNA kit), a 3 nt UMI (unique molecular identifier), a 6 nt unique barcode and half a NlaIII digestion site (CA) such that NlaIII cutting site is reconstituted upon self-ligation of adapters (CATG). The barcodes were designed with a hamming distance of two. Bottom sequences contained a phosphorylation site at the 5’ end. Adapters were produced as standard desalted primers. Top and bottom sequences were annealed at a 1:1 ratio in annealing buffer (10 mM Tris pH 7.5–8.0, 50 mM NaCl, 1 mM EDTA) by immersing tubes in boiling water, then let to cool to room temperature. The oligo sequences can be found in Supplementary Table 1.

### CEL-seq primers

The RT primer was designed according to the Yanai protocol^4^ with an anchored polyT, a 8nt unique barcode, a 6nt UMI (unique molecular identifier), the 5’ Illumina adapter (as used in the Illumina small RNA kit) and a T7 promoter. The barcodes were designed such that each pair is different by at least two nucleotides, so that a single sequencing error will not produce the wrong barcode. Primers are desalted at the lowest possible scale, stock solution 1 µg/µL. The oligo sequences can be found in Supplementary Table 2.

### Raw data preprocessing

First mates in the raw read pairs (i.e. “R1” or “read1”) conform to a layout of either:

5′-[3 nt UMI][8 nt barcode]CA[gDNA]-3′

in the case of gDNA (DamID and AluI restriction) reads, or

5′-[6 nt UMI][8 nt barcode][unalignable sequence]-3′

in the case of transcriptomic reads.

In the case of transcriptomic reads, the second mate in the read pair contains mRNA sequence.

Raw reads were processed by demultiplexing on barcodes (simultaneously using the DamID and transcriptomic barcodes), allowing no mismatches. The UMI sequences were extracted and stored alongside the names of the reads for downstream processing.

### Sequence alignments

After demultiplexing of the read pairs using the first mate and removal of the UMI and barcode sequences, the reads were aligned. In the case of gDNA-derived reads, a ‘GA’ dinucleotide was prepended to the sequences of read1 (‘AG’ in the case of AluI), and read1 was then aligned to a reference genome using bowtie2 (v.2.3.2) using parameters --seed 42 --very-sensitive -N 1. For transcriptome-derived reads, read2 was aligned using tophat2 (v2.1.1) using parameters --segment-length 22 --read-mismatches 4 -- read-edit-dist 4 --min-anchor 6 --min-intron-length 25 --max-intron-length 25000 --no-novel-juncs --no-novel-indels --no-coverage-search --b2-very-sensitive --b2-N 1 --b2-gbar 200 and using transcriptome-guiding (options --GTF and --transcriptome-index). Human data was aligned to hg19 (GRCh37) including the mitochondrial genome, the sex chromosomes and unassembled contigs. Transcriptomic reads were aligned by making additional use of transcript coordinates obtained from GENCODE (v26) https://www.gencodegenes.org/releases/grch37_mapped_releases.html supplemented with ERCC mRNA spike-in sequences https://assets.thermofisher.com/TFS-Assets/LSG/manuals/cms_095047.txt. mESC data was aligned to reference genomes generated by imputing 129S1/SvImJ and CAST/EiJ SNPs obtained from the Sanger Mouse Genomes project [http://www.sanger.ac.uk/science/data/mouse-genomes-project^5^, onto the mm10 reference genome. The mitochondrial genome, sex chromosome and unassembled contigs were used in the alignments. Transcriptomic reads were aligned using a GTF file with transcript annotations obtained from ENSEMBL (release 89) [ftp://ftp.ensembl.org/pub/release-89/gtf/mus_musculus/Mus_musculus.GRCm38.89.gtf.gz]. Both human and mouse references were supplemented with ERCC mRNA spike-in sequences [https://assets.thermofisher.com/TFS-Assets/LSG/manuals/cms_095047.txt]. For both genomic and transcriptomic data, reads that yielded an alignment with mapping quality (BAM field ‘MAPQ’) lower than 10 were discarded. For the genomic data, reads not aligning exactly at the expected position (5′ of the motif, either GATC in the case of DpnI restriction, or AGCT in the case of AluI restriction) were discarded. For the transcriptomic data, reads not aligning to an exon of a single gene (unambiguously) were discarded. The mESC reads were assigned to the 129S1/SvImJ or CAST/EiJ genotype by aligning reads to both references. Reads that align with lower edit-distance (SAM tag ‘NM’) or higher alignment-score (SAM tag ‘AS’) in case of equal edit-distance to one of the genotypes were assigned to that genotypes. Reads that aligned with equal scores to both genotypes were considered of ‘ambiguous’ genotype.

### PCR duplicate filtering

For the genomic data (DamID and AluI-WGS), the number of reads per motif, strand and UMI were counted. Read counts were collapsed using the UMIs (i.e. multiple reads with the same UMI count as 1) after an iterative filtering step where the most abundant UMI causes every other UMI sequence with a Hamming-distance of 1 to be filtered out. E.g, observing the three UMIs ‘AAA’, ‘GCG’ and ‘AAT’ in decreasing order would count as 2 unique events (with UMIs ‘AAA’ and ‘GCG’, since ‘AAT’ is within 1 Hamming distance from ‘AAA’). For the data from KBM7 (a near-complete haploid cell line) at most 1 unique event per motif and strand was kept. For the mESC data at most 1 unique event per motif, strand and genotype was kept, or 2 unique events, if the genotype of the reads at that position could not be resolved.

### Filtering of samples

Only single-cell samples with at least 10^3.7^ unique DamID events or at least 10^3^ unique transcripts were taken into consideration for the analyses. These cutoffs were applied jointly for analyses where both genomic and transcriptomic signals were used.

### Binning and calculation of OE values

DamID and WGS data was binned using non-overlapping bins. Binsizes were 100kbp for untethered Dam and 500kbp for Dam-LmnB1 DamID data, 100kbp for WGS data and 500kbp for all hybrid mESC data where genotype-specific counts were used. For analyses at TSS and CTCF sites, binsizes were 10bp. In order to calculate observed-over-expected (OE) values, the mappability of each motif (GATC or AGCT) was determined by generating 65 nt. long sequences (in both orientations) from the reference genome(s) and aligning and processing them identically to the data. By binning the *in-silico* generated reads, the maximum amount of mappable unique events per bin was determined. OE values were calculated using

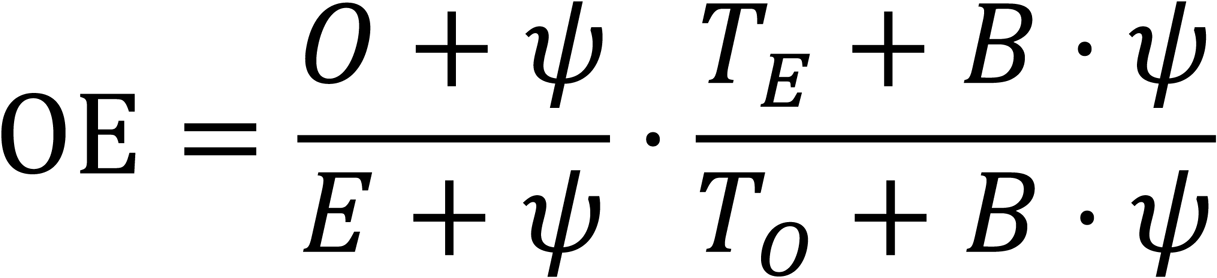

where *O* is the number of observed unique methylation events per bin, *E* is the number of mappable unique events per bin, *ψ* is the pseudocount (1, unless otherwise stated), *T*_*O*_ and *T*_*E*_ are the total number of unique methylation events observed cq. mappable in the sample and *B* is the number of bins. For analysis across multiple windows, e.g. windows around TSSs or CTCF sites, *O* and *E* are summed across the windows, prior to calculation of the OE values. For the definition of “contact”, regions with OE values >= 1 were considered as “in-contact”. For further details and justification, see Kind et al., 2015^1^ and FigS2A in particular.

### H3K4me3, H3K36me3 and DNase data (external datasets)

H3K4me3, H3K36me3 and DNase data was obtained from ENCODE (GSM788087, GSM733714 and GSE90334_ENCFF038VUM, respectively) as processed bigWig files. In order to calculate OE values for these datasets, whole-genome mappability as determined by the ENCODE project was used (wgEncodeCrgMapabilityAlign36mer).

### Independent transcription dataset

For Fig2G independent expression data was used from GSE56465. (only KBM7 haploid samples).

### Untethered Dam enrichment at TSSs and CTCF sites

For the analyses at TSSs, one isoform per gene was chosen from the gene annotations, by taking preferentially isoforms that carry the GENCODE “basic” tag, have a valid, annotated CDS (start and stop codon, and CDS length that is a multiple of 3nt.), and ties are broken by the isoform with longest CDS, and shortest gene length (distance from first to last exon). As TSS, the most 5′ position of the first exon was taken. CTCF sites were obtained by integrating ENCODE ChIPseq data (wgEncodeRegTfbsCellsV3, K562 CTCF ChIPseq tracks from GSE30263) with CTCF motif sites (factorbookMotifPos obtained via the UCSC genome browser^6^). Only CTCF ChIPseq peaks that contained a CTCF binding motif with score of at least 1.0 within 500nt. of the center of the ChIPseq peak were considered. The ChIPseq peaks were subdivided by ChIPseq binding score, and the group of peaks with maximum score (of 1000) was subdivided into two groups by the motif score, such that 4 approximately equal-sized groups of CTCF-bound loci were obtained.

### logFC between contact/no contact groups of samples

logFCs between single-cell samples that showed contact and those that show no contact (see Fig3A) was performed as follows: In bins across the genome (500kb. for Dam-LmnB1, 100kb. for untethered Dam) the logFC in expression was calculated between samples that have a DamID OE value ≥ 1 vs. samples that have a DamID OE value lower than 1, for every bin that has (1) at least 103.4 mappable GATCs per 100kb and (2) contains at least 3 single-cell samples per group and (3) has a mean transcriptional level of at least 10 RPM across all single-cell samples. Comparison scDamID&T to Kind Cell 2015 data. For the comparisons with individual measurements of single-cell DamID and single-cell transcriptomics (CELseq) with scID&T in Fig1 the scID&T data was made comparable to the published data by (1) truncating the reads at the 3′ end such that after barcode (and in the case of scDamID adapters) removal the same number of nt. of gDNA is remaining. Furthermore, UMIs were completely left out of the consideration for the DamID measurements, and for the transcriptional measurements, the UMIs were truncated to 4nt. to make the data comparable to the published CELseq data. The data were obtained from GSE69423.

By figure details on the statistics can be found in Supplementary Table 3. All computational codes used for this study are available upon request.

