## Supplementary figures and images for "Simultaneous quantification of protein-DNA contacts and transcriptomes in single cells"

### Supplemental Figures

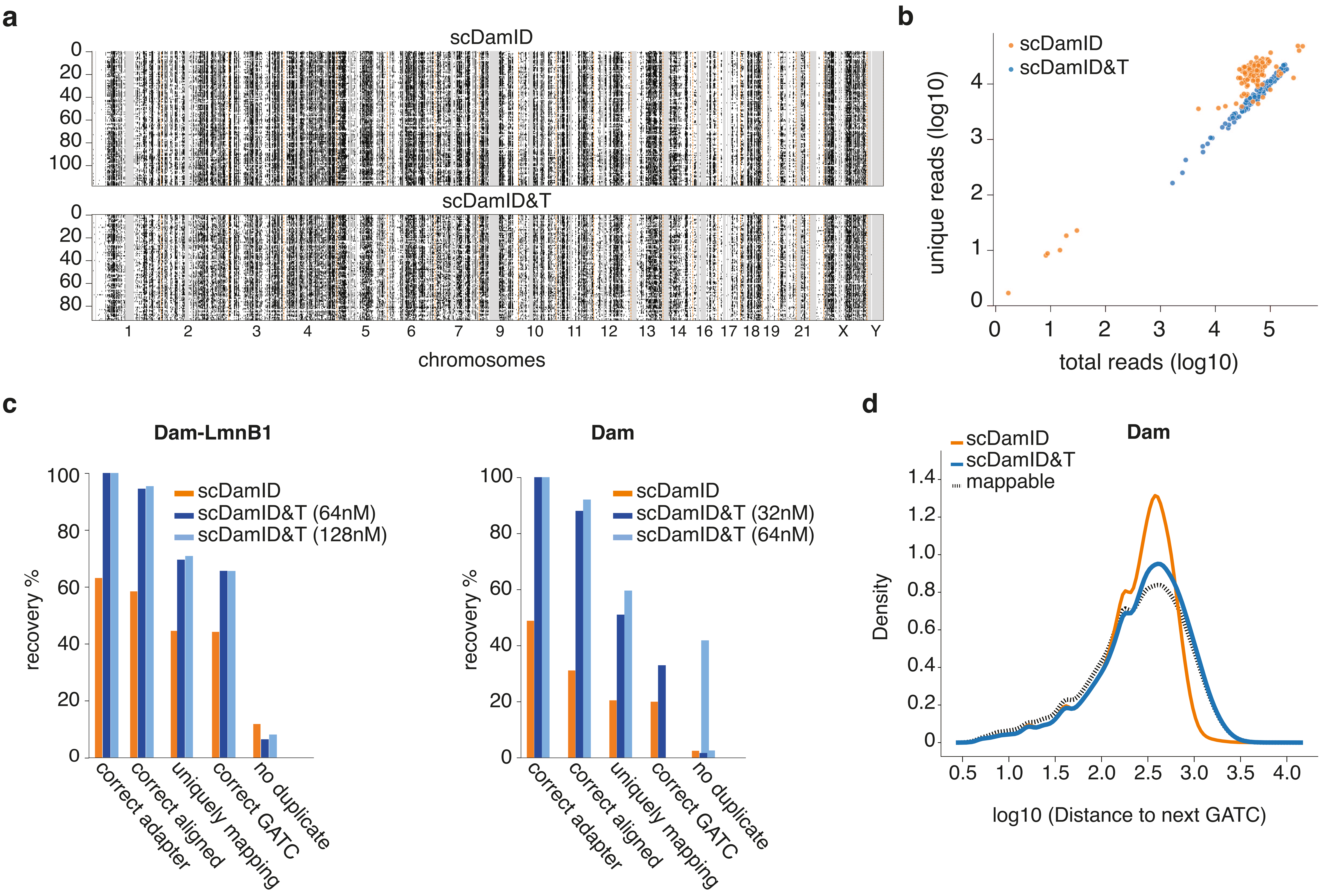

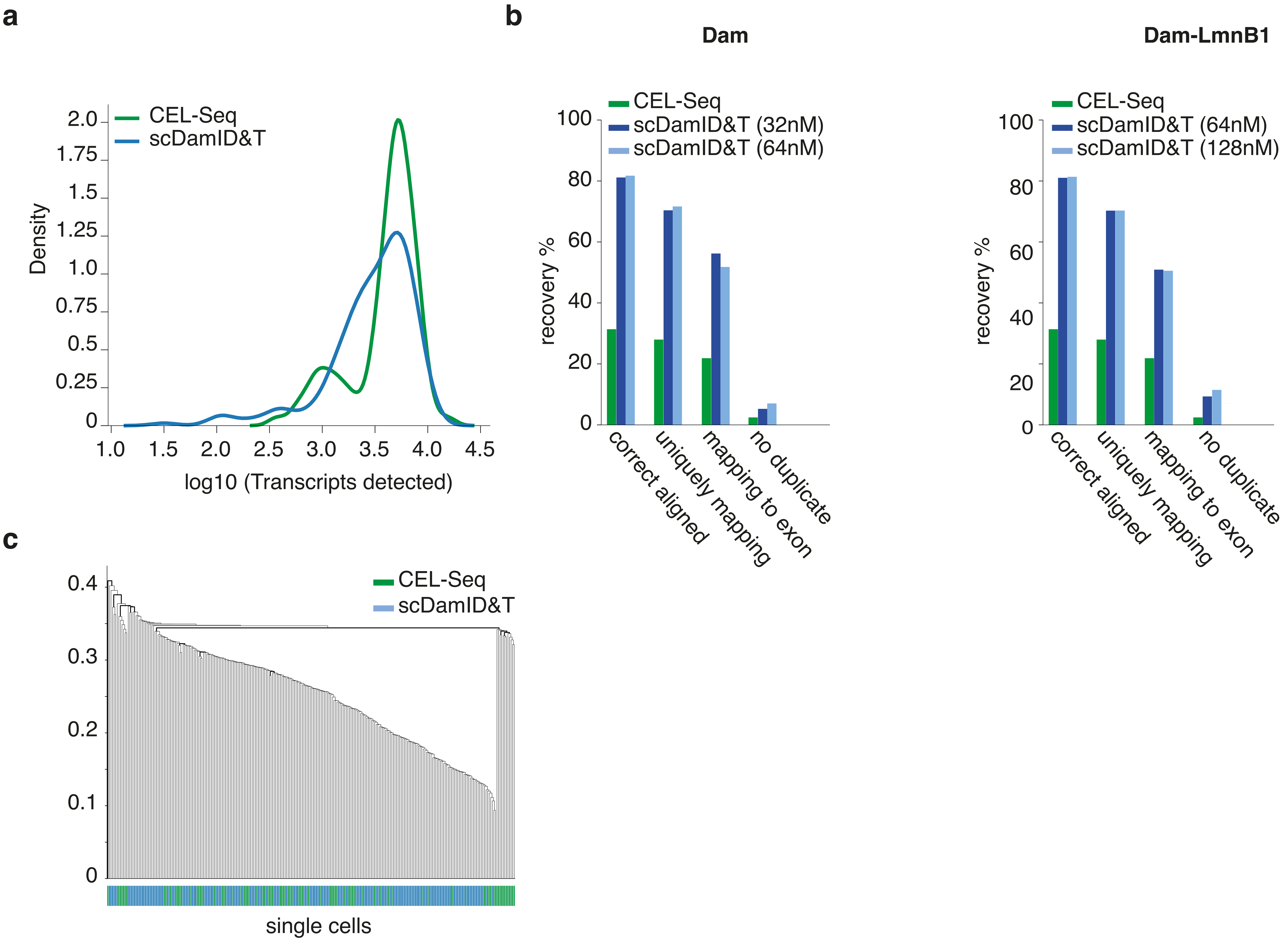

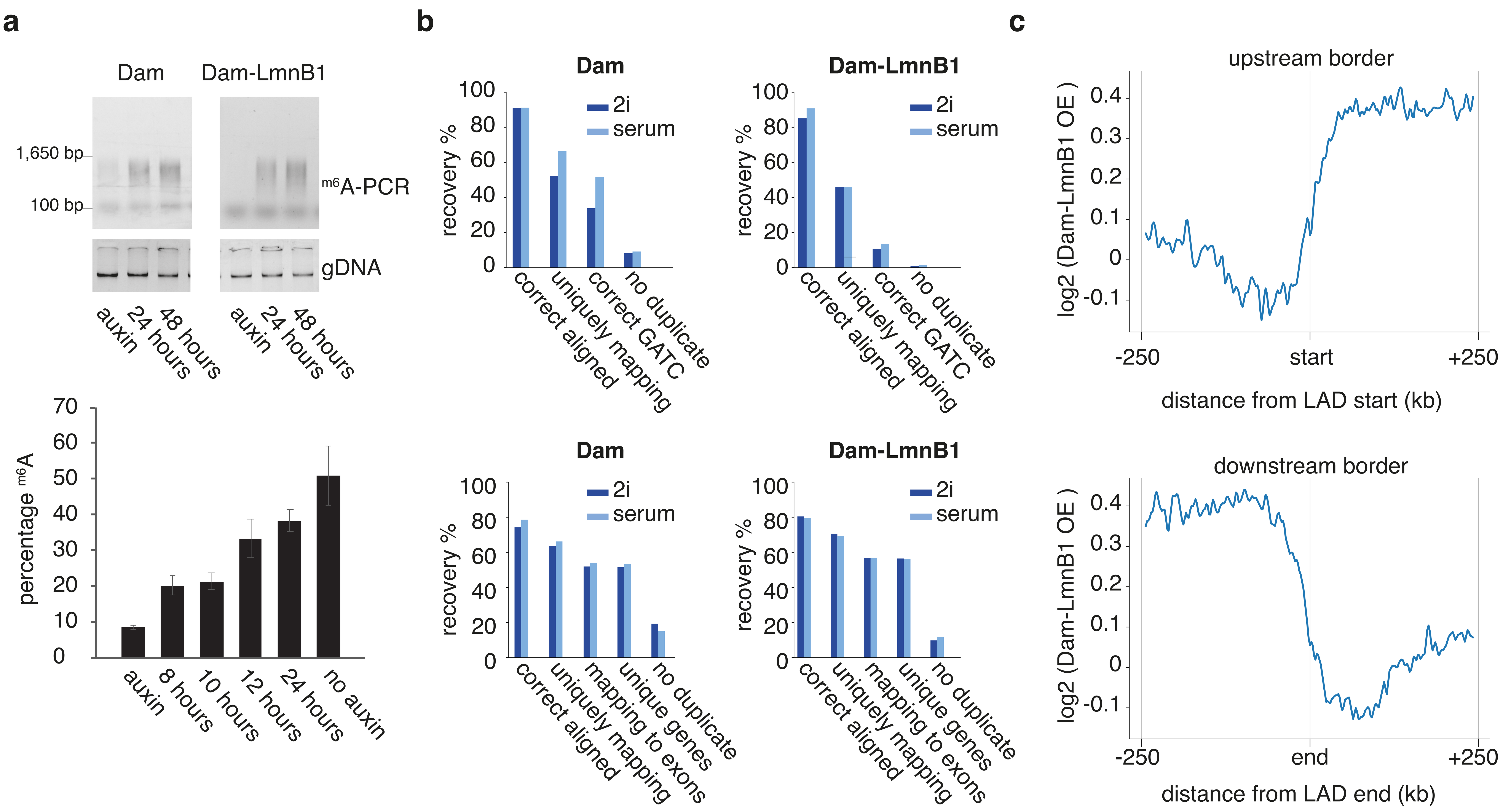

#### Figure S4

**a**

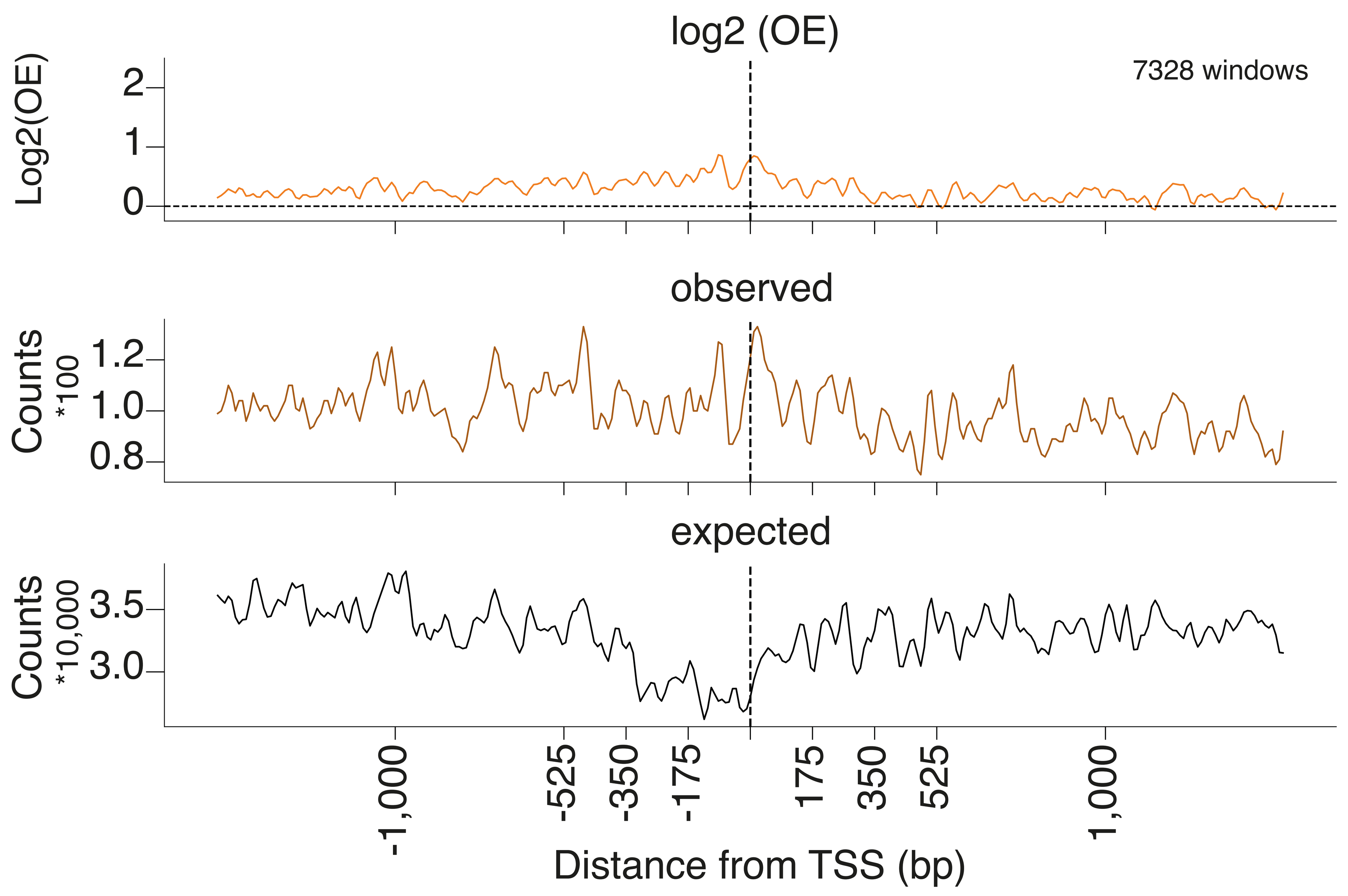


**b**

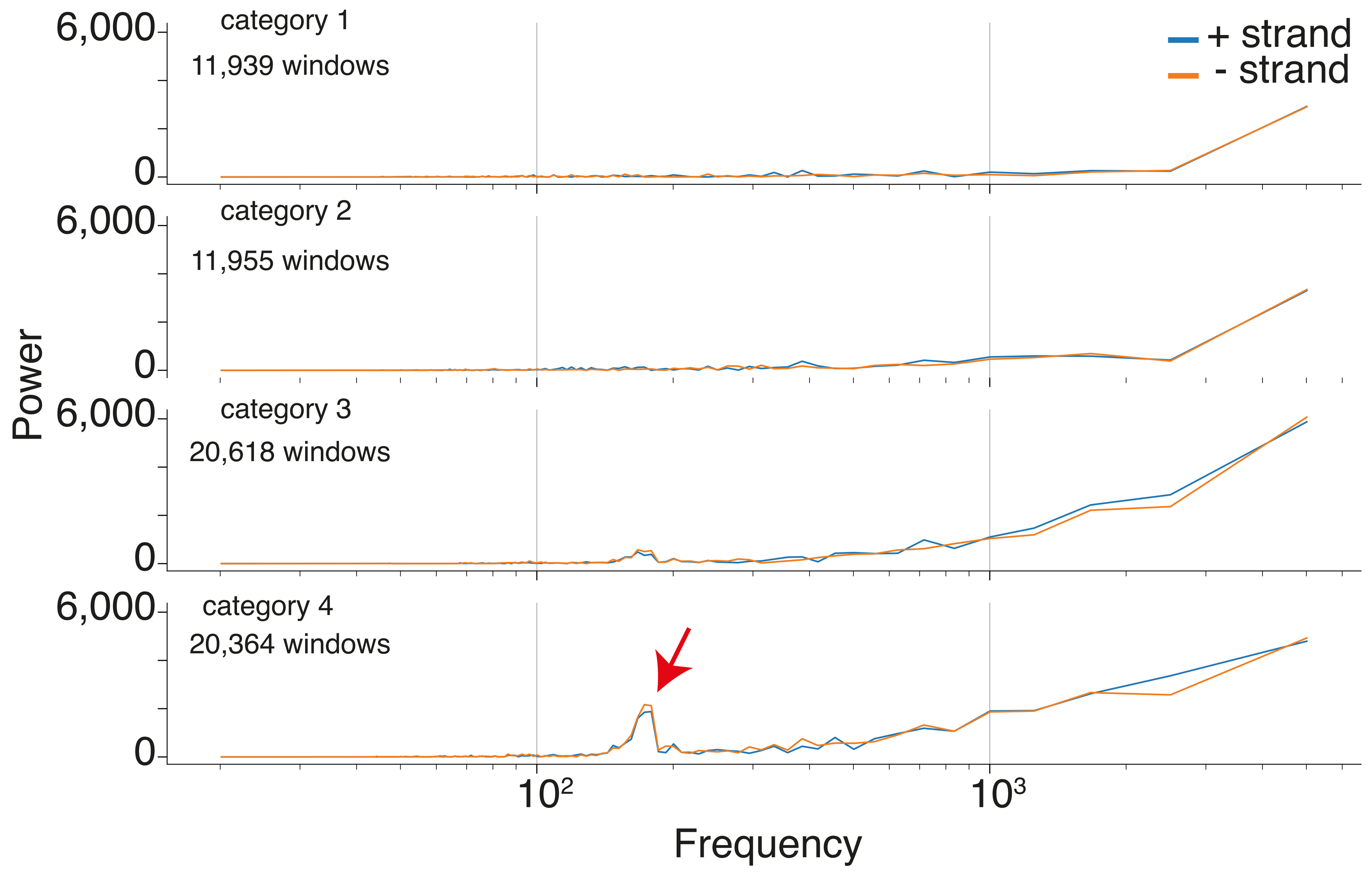


**c**

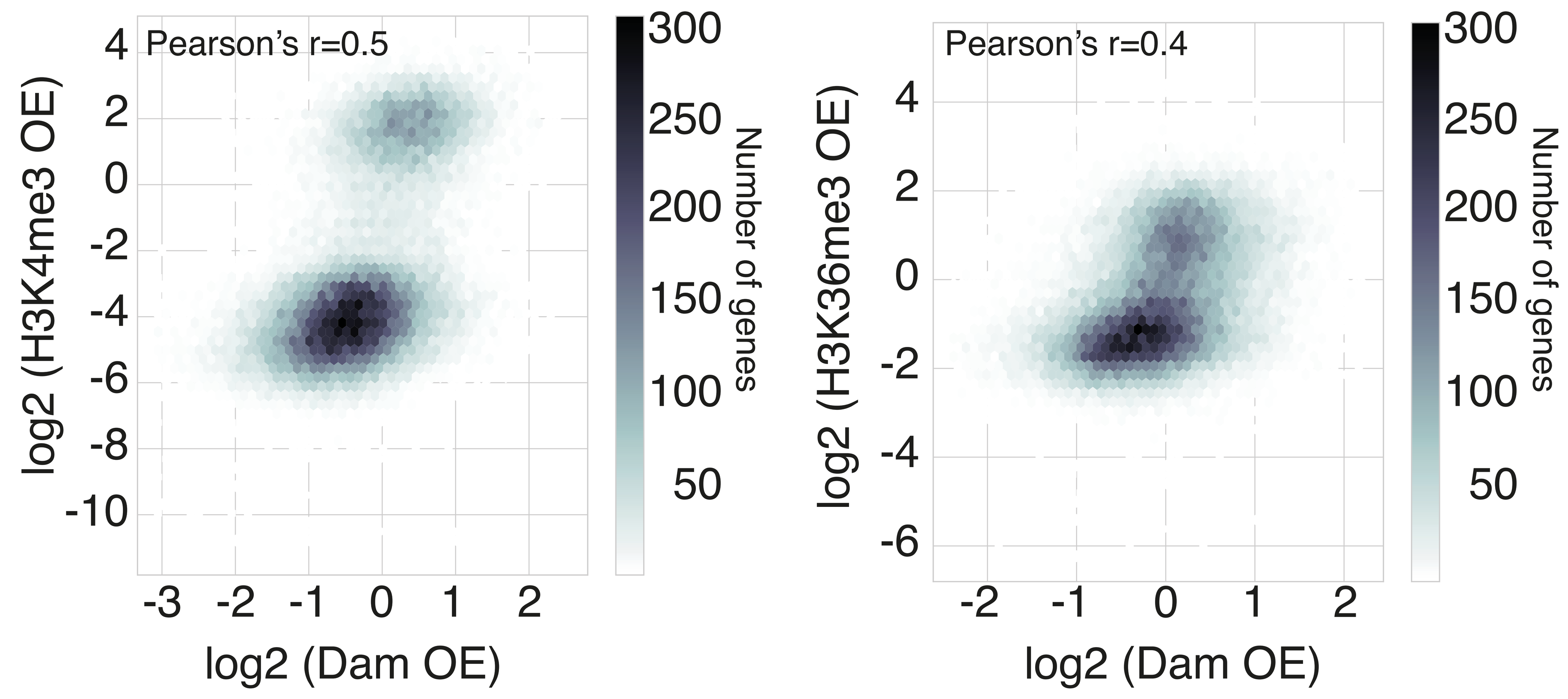

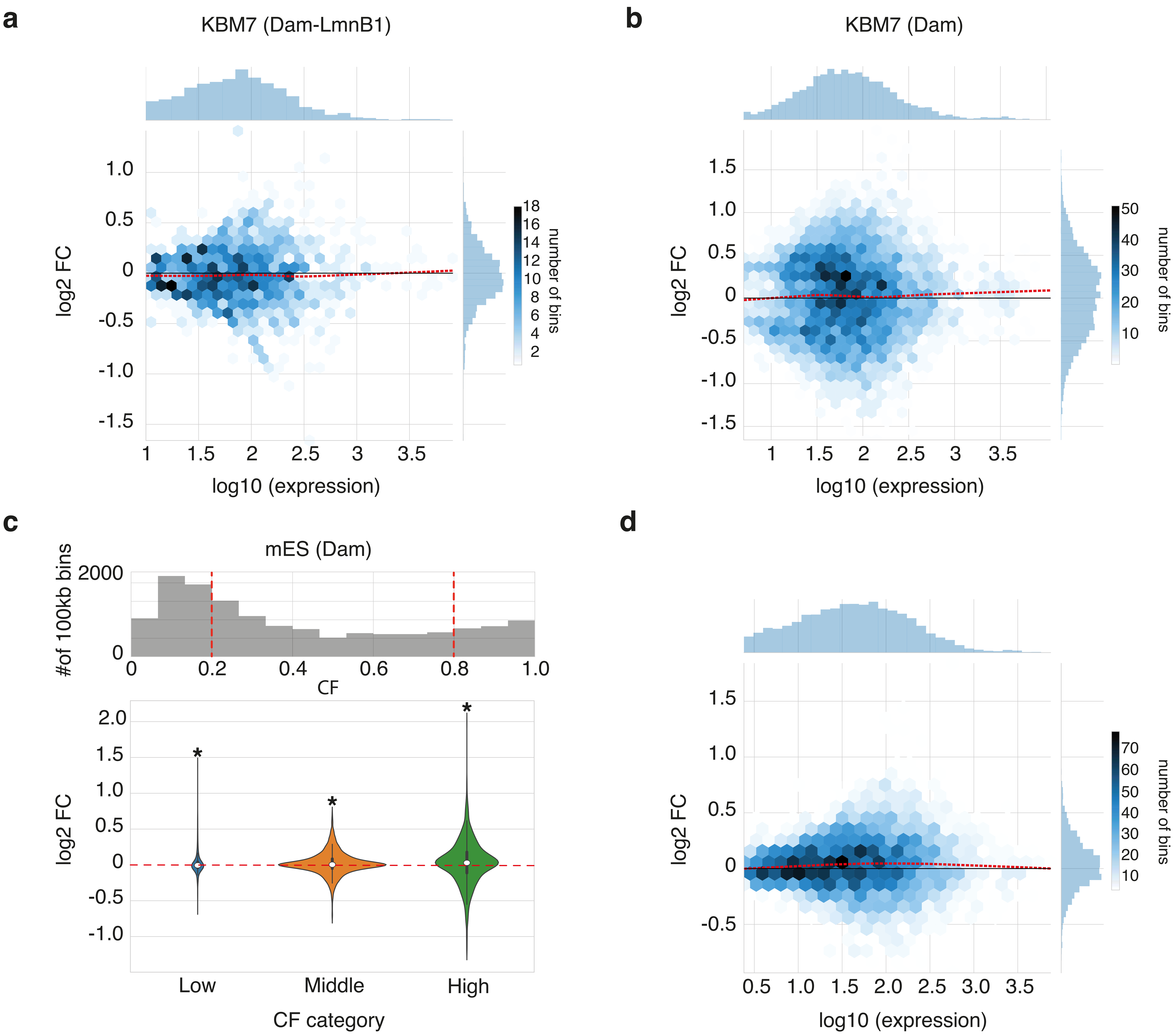

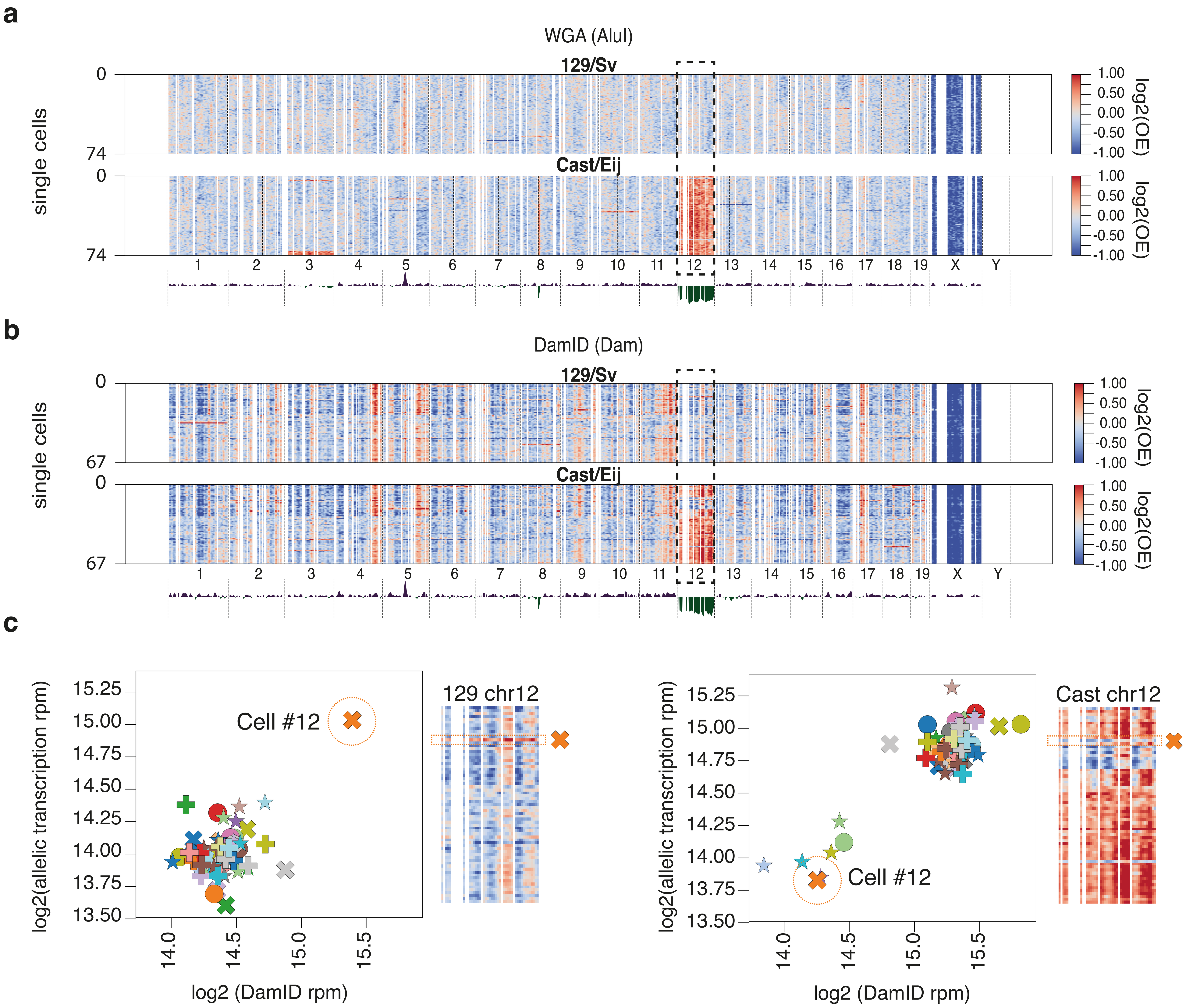

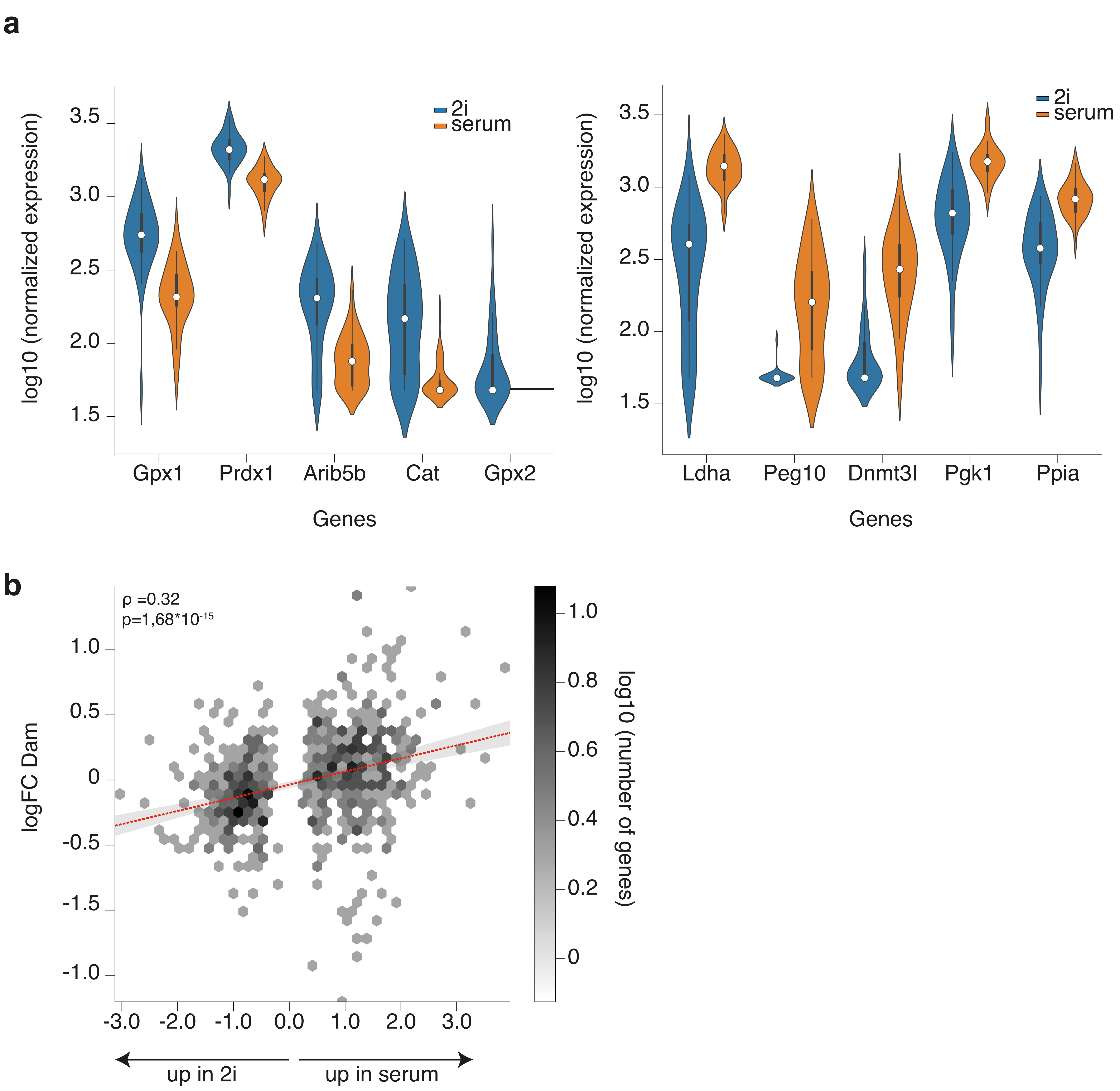
